# Comprehensive non-black box classification of highly correlated ecological time series pairs containing many zeros: the case of gut microbiome of mice

**DOI:** 10.1101/2024.03.15.585153

**Authors:** Rie Maskawa, Hideki Takayasu, Tanzila Islam, Lena Takayasu, Rina Kurokawa, Hiroaki Masuoka, Wataru Suda, Misako Takayasu

## Abstract

We developed a new data analysis method, named Coexistence–Exclusion–Synchronization– Antisynchronization (CESA), to reveal statistically significant correlations from a set of integer compositional abundance time series of Operational Taxonomic Unit (OTU) data of mouse gut microbiota. First, time series are transformed to 0 (absence) and 1 (presence), and statistical tests are applied to extract significant coexistence and mutual exclusion relationships. Subsequently, for all pairs, the difference time series are transformed to +1 (up), 0 (even), and −1 (down), and synchronized and antisynchronized pairs are classified based on statistical tests after carefully removing the effect of spurious correlation caused by changes in compositional shares. We performed a comprehensive classification of all pairs based on the p-values in terms of coexistence and synchronization, including time series data with many zeros, which are difficult to analyze using conventional methods. We found that almost all OTUs (419 out of 420) have significant correlations with at least one OTU in one of the four characteristics: coexisting, exclusive, synchronizing, or antisynchronizing. Considering OTU pairs, about 25% of all possible pairs (22,356 out of 87,990) show a high correlation with the p-values less than 10^-5^, which is less than the inverse of the total number of pairs. Interaction among phyla are summarized as a network diagram.

**Author summary:** The gut microbiota ecosystem is often thought to be stable. However, when observed over a long period, there are turnovers in the microbiota, each OTU time series is highly non-stationary, and even species with high overall abundance are often observed to have zero values in some periods. In this study, we developed a comprehensive data analysis method for extracting significant correlations between any pair of OTUs, including OTUs whose observed values contain many zeros or exhibit clear non-stationarity, for which processing methods have not yet been established. We focused on pairwise correlations in terms of coexistence, exclusivity, synchrony, and antisynchrony of increase/decrease, and all combinations of pairs were checked by statistical tests based on the p-values. In order to remove spurious correlations in compositional time series, a new method was introduced to correct the sample sizes for the remaining OTUs, hypothetically assuming a situation in which one OTU was not present.

Low abundance OTUs are often overlooked in traditional analyses. However, it becomes evident that all OTUs, including those with low abundance, interact strongly with each other. Additionally, our findings suggest that coexistence and synchrony can be summarized as cooperative relationships, while exclusion and antisynchrony can be summarized as antagonistic relationships. Cooperative interactions are more likely to occur between pairs of OTUs in the same phyla, and antagonistic interactions are more likely to appear between OTUs in different phyla. The time series data analysis method developed in this paper includes no black-box, making it broadly applicable to compositional time series data with integer values.

## Introduction

The gut microbiome is a highly complex and dynamic ecosystem that has attracted considerable scientific interest due to its profound influence on host health and development [1–6]. This complex microbial community orchestrates a variety of physiological processes within the host, ranging from nutrient metabolism to modulation of the immune system. To unravel the full extent of these effects and their temporal dynamics, the study of the mouse gut microbiome has become increasingly important. The gut microbiome is a vast collection of microorganisms that reside in the gastrointestinal tract. It includes bacteria, viruses, fungi, and other microbes that form a dynamic and complex ecosystem. This ecosystem evolves over time, with microbial composition and functions adapting in response to various factors, including diet, age, and environmental exposures. Understanding the temporal dynamics of the gut microbiome is critical because it can shed light on the mechanisms underlying health and disease. Recent advances in high-throughput sequencing technologies have revolutionized our ability to characterize the gut microbiome. Numerous studies have investigated its composition and functions in various contexts, revealing its role in metabolic disorders, immune system regulation, and even neurological diseases [7–11].

From a data analysis perspective, time series data of bacterial flora ecosystems are challenging to analyze for various reasons, and numerous methods of data analysis have been developed [12]. In most cases, the total number of abundances observed in a single measurement is fixed, thus, the abundances are relative rather than absolute quantities. Therefore, it needs to be considered that spurious correlations may arise, wherein an increase in the number of one species leads to a decrease in the number of other species [13,14]. In a typical bacterial ecosystem, approximately 1000 OTUs are observed, and their abundance varies by more than three orders of magnitude, with high- and low-abundance species exhibiting markedly different behaviors [15,16].

For time series of high abundance OTUs, the z-score transformation [17] and the log-ratio transformations [18,19] are standard pretreatments. Among various proposed models, the stochastic logistic model is known to successfully reproduce the basic characteristics of high-abundance stationary time series [15,20,21]. To estimate mutual interactions between species, the Lotka–Voltera equation is used as a basic dynamical equation [22,23]. The analysis of linear and nonlinear correlations [24,25] and network analyses [26] is also under intensive investigation. Approaches to estimating causal relationships between species from time series data are also being explored [27,28].

Thus, data analysis methods for high-frequency and stationary cases are well-developed, whereas those for low-frequency time series and clearly non-stationary cases are still under development. Time series of low-abundance species contain many zeros, and the analysis of such data requires different processing than that of high-frequency cases. For example, the log-ratio method cannot be directly applied to data containing zeros. For time series with many zeros, studies have been conducted since the 1960s to examine correlations between species by binarizing them as present or absent [29]. It is reported that Pearson’s correlation coefficient, the most fundamental quantity characterizing correlations, is biased toward a positive value in the case of binary data depending on the proportion of zeros included, and it is argued that a null model based on a hypergeometric distribution should be considered [30]. Recently, a machine learning approach was introduced for time series with many zeros to infer the interaction network between bacterial species based on the hurdle model. This approach applies the group lasso penalty to the matrix of time series data, which is converted into binary form (1 for presence and 0 for absence) [31].

In this paper, we introduce a novel non-black-box data analysis method named Coexistence– Exclusion–Synchronization–Antisynchronization (CESA) using the OTU time series in the gut microbiota of seven mice. This allowed us to comprehensively reveal significant correlations among all OTU pairs, including low-abundance or non-stationary cases, with a new framework for adjusting spurious correlations caused by compositional property. The paper is organized as follows, with a flow-chart shown in Fig 1. In the next Results section, we first describe the data we are dealing with. We examine the distribution of frequencies per OTU as a fundamental property of the data, focusing on all pairs of 420 OTUs common to all mice, totaling 87,990 pairs. After binarizing the time series data in terms of presence, we test for each pair of time series whether there is a significant coexisting or exclusively non-coexisting relationship by calculating the p-value based on the hypergeometric distribution, the Fisher exact test. Next, focusing on the change in the time series, the data are transformed into ternary values, +1 for an increase, 0 for no change, and −1 for a decrease, and pairs with significant synchronous and antisynchronous relationships are extracted through a statistical test after removing spurious correction caused by the compositional nature of data. In the Discussion section, we address the validity of the classified correlations extracted in this way. In the Materials and Methods section, we introduce details of our new correction method of removing spurious correlations of synchronization. Additionally, details about the mathematical formulations used in this paper are explained in this section.

**Fig 1.**
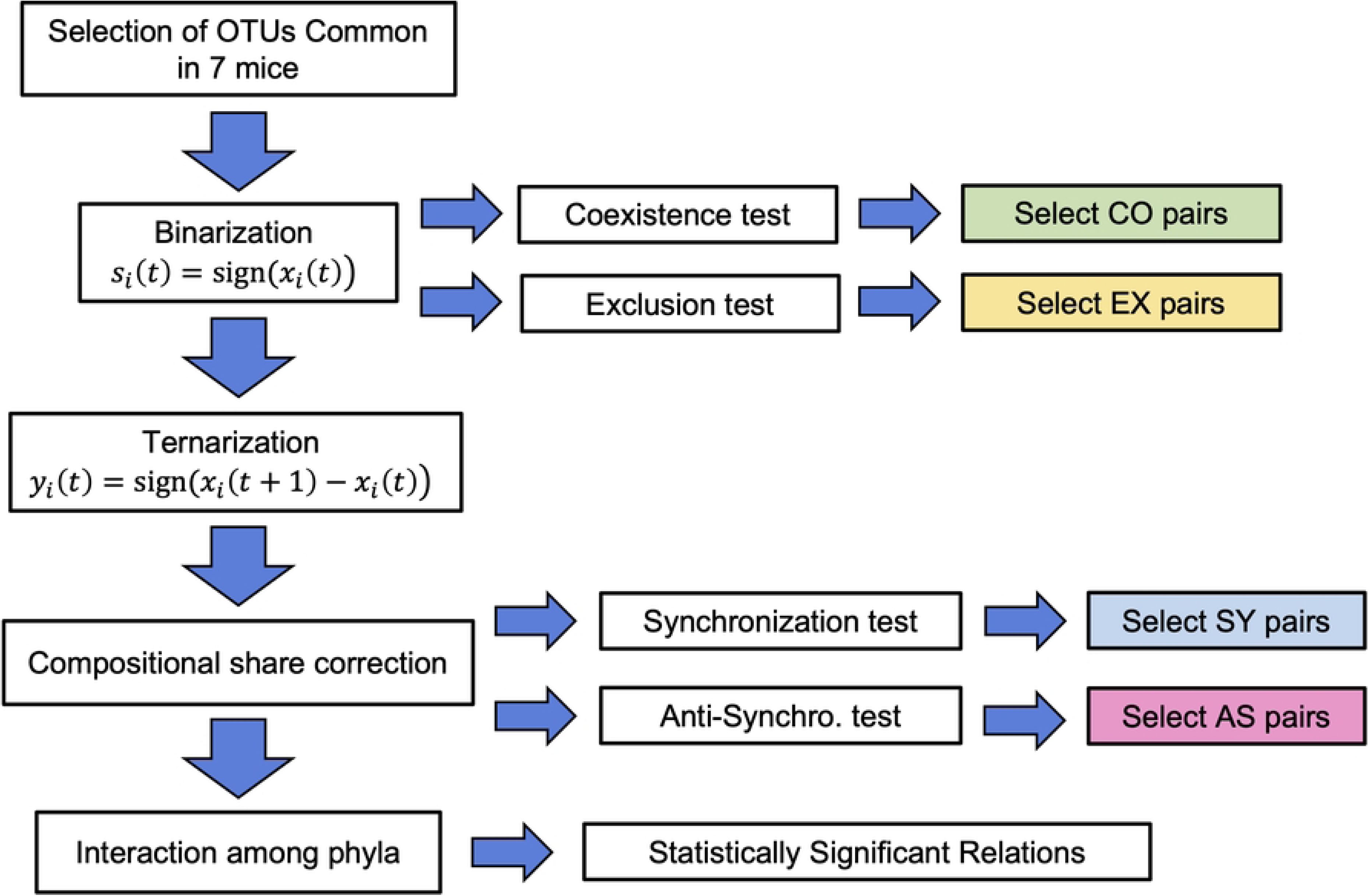
Flowchart of the data analysis methods in this paper. For any two OTUs, four correlations are extracted based on the p-values: CO, EX, SY, and AS. Finally, the OTUs are grouped into phyla and then analyzed for inter-phyla correlations.

Our new method is simple, does not involve a black box, and is versatile enough to be immediately applied to any similar time series data set.

## Results

### The data and basic properties of the time series

In this study, we use the time series data of the mouse gut microbiome to demonstrate the broad applicability of our method. Specifically, our analysis focuses on the OTUs within seven mice (designated M1, M2, M3, M4, M5, M7 and M8) over their entire lifespan from birth to natural death. The average lifespan of these mice is 923 days, with individual lifespans ranging from 827 to 1,044 days [32]. An OTU represents a group of closely related microorganisms, typically observed through gene sequencing, specified by a natural number up to five digits, such as OTU279. In one observation, 3,000 samples are detected from fecal microbiomes, demonstrating the presence of a specific set of genetic information across different samples. The average observation interval was 4.3 days, resulting in about 200 data points for each mouse. This dataset serves as a critical component of our research, allowing us to examine the temporal variation in gut microbiome composition in different mice.

The total number of OTUs in the data for all seven mice over their lifetime is approximately 4 million, consisting of 1,230 OTUs. Fig 2a shows the cumulative distribution of the sampling probability for each OTU in our entire data on a log–log scale. It is a typical fat-tailed distribution with a range of variation that is spread out over five digits. This distribution can be approximated by the well-known log-normal distribution (red curve), which is consistent with the stochastic logistic model [15].

**Fig 2.**
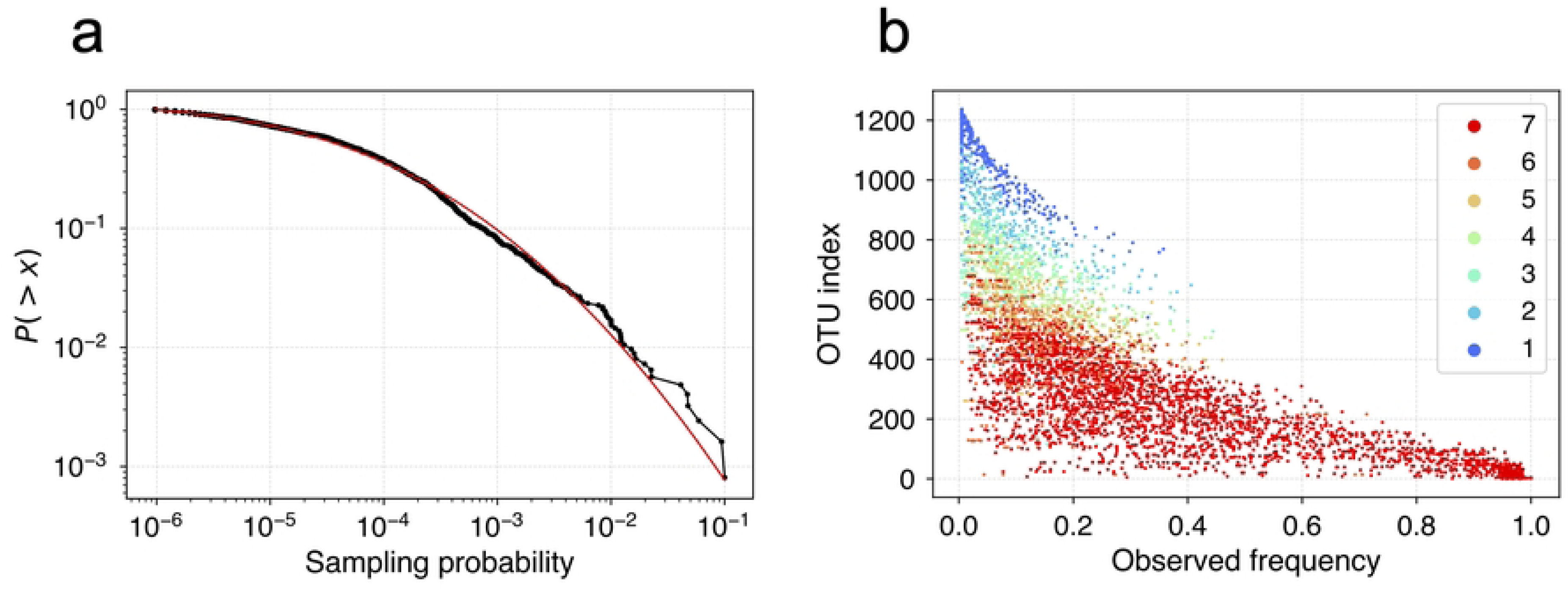
Basic properties for all OTUs. Fig 2a. Complementary cumulative distribution of sampling probabilities for all 1230 OTUs observed in 7 mice plotted in a log-log plot. The red curve shows a log-normal distribution fitted with μ=−10.1 and σ=2.47. Fig 2b. Observed percentage of non-zeros plotted for all OTUs for all mice individually. The vertical axis of the OTU index is numbered in the descending order of sampling probability. The color indicates the number of mice observed: red for seven mice, orange for six mice, …, blue dots indicate the OTU appearing only in 1 mouse. The number of OTUs common to all seven mice is 420.

In Fig 2b, the percentage of non-zeros observed in each OTU time series is plotted for each mouse. The vertical axis is the OTU index, sorted in descending order of sampling probability from Fig 2a. The colors indicate the number of mice on which the OTU was observed. For example, there are seven red dots for seven mice of the same OTU index plotted horizontally, six orange dots for six mice, etc., and the OTU of the blue dots is observed in only one mouse. This plot shows that the percentage of 0 is generally high, even for popular OTUs that are common to all mice. It is also confirmed that there is no OTU that is always present in all observations. This result would indicate that even a highly abundant OTU is not stationary throughout life. The number of OTUs common to all seven mice shown as red dots in Fig 2b is 420, and in the following analysis we will focus on all combinations of pairs of these common OTUs, 87,990 pairs.

### Detection of coexistence and exclusion pairs

As described in the previous section, most of the time series in our data contain many zeroes. We were interested in the correlation of presence and absence between the time series of two different OTUs of each mouse. For this purpose, we transformed all time series {*x_i_*(*t*)} into a binary form (1 for presence and 0 for absence), where “sign” designates the sign function (1 for positive, 0 for 0, and −1 for negative), as follows:

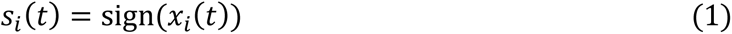

Focusing on the binary time series of OTU-*i* and OTU-*j* of the *k*-th mouse, there are four cases: (1,1), (1,0), (0,1) or (0,0). At each observation time step, and we count the numbers. Let *a* be the number of co-presence (1,1), *b* the number of (1,0), *c* the number of (0,1), and *d* the number of (0,0), which is the case of co-absence; the sum of these numbers makes the length of the time series, *T*=*a+b+c+d* (see Materials and Methods for mathematical formulation).

As a null model, we assume randomly shuffled time series for OTU-*i* and OTU-*j*, where the numbers 0 and 1 are conserved for each OTU. If the actual number of *a* is larger than the random case, it means that OTU-*i* and OTU-*j* tend to coexist; if *a* is smaller than the random case, it means that these OTUs tend to be exclusive. It is known that the distribution of {*a, b, c, d*} follows a hypergeometric distribution [30], thus, we can estimate the corresponding p-value by using Fisher’s exact test for a 2×2 contingency table. As the degree of freedom of this contingency table is 1, the p-value for coexistence is calculated by summing the probability of the null model where the number of (1,1) is equal to or greater than *a*. This value can be small in the case of *ad-bc>0*. Similarly, the p-value for exclusion is calculated by summing the probability of the null model where the number of (1,1) is equal to or smaller than *a*. This value can be small in the case of *ad-bc*<0. It should be noted that if one of the OTU is always present, then the p-value is automatically 1 for both coexistence and exclusion, because the set of numbers {*a, b, c, d*} of any randomly shuffled time series is identical to the real data. In fact, to make the p-value small, both OTU should have a certain amount of zeros for both coexistence and exclusion.

Fig 3a shows the cumulative distributions of p-values of one mouse (M1) for coexistence (green line) and for exclusion (orange line) compared to the theoretical distribution of the p-values of the null model (gray line), which follows the uniform distribution. We found that it is reasonable to set a threshold of statistical significance for the p-value at 10^-5^ (gray dotted line) since this probability is smaller than the inverse of the total number of observed OTU pairs, 1/87990≒1.1×10^-5^. Fig 3b displays the resulting scatter plot for all combinations of OTUs of mouse M1. The horizontal axis is the number of coexistences (*a*), and the vertical axis is *(ad-bc)/T^2^*, a value characterizing the strength of coexistence normalized by the length of the time series. The green and orange dots indicate statistically significant OTU pairs of coexistence *(ad-bc>0)* and exclusion *(ad-bc<0)*, respectively, while the gray dots indicate non-significant cases. In this instance, approximately 7% of OTU pairs show significance in coexistence, and approximately 1% are significant in exclusion. As observed in the figure, the values of a are always small in cases of significant exclusion, while in cases of coexistence, the values of a are widely distributed.

**Fig 3.**
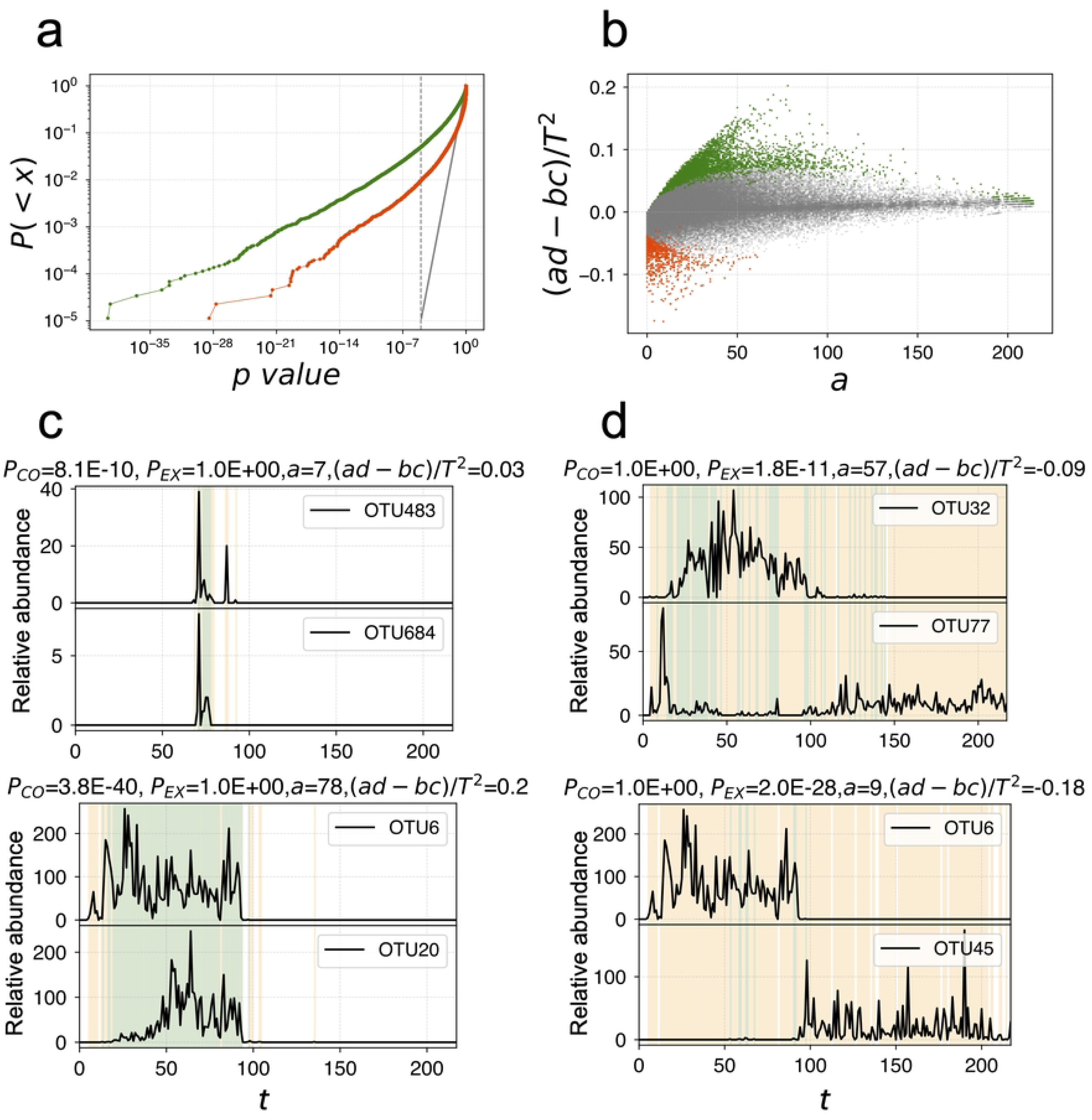
Coexistence and exclusion for one mouse. Fig 3a. Cumulative distribution of the p-values for coexistence (green) and exclusion (orange) for all combinations of OTUs in the log-log plot. The distribution of p-values in the case of the null model follows the uniform distribution for both cases (gray). The dashed line indicates the statistical significance threshold, 10^-5^. Fig 3b. Scatter plot of the values *a* (number of coexistence) vs *(ad-bc)/T^2^*. Green and orange dots show significant cases of coexistence and exclusion, respectively, and gray dots are plotted for non-significant cases. Fig 3c. Examples of OTU pairs of time series for coexistence. Green and orange shades show coexistence (1,1) and exclusion (1,0) or (0,1), respectively, while (0,0) are in white. Top: The probability of appearance is very low for both, but the timing of appearance tends to coincide. Bottom: Abundances are high for these OTUs, and both disappear in the second half of life. Fig 3d. Examples of OTU pairs of time series for exclusion. Green and orange shades show coexistence (1,1) and exclusion (1,0) or (0,1), respectively, while (0,0) are in white. Top: OTU6 disappears in the second half of life, while OTU45 appears in the second half of life. Bottom: OTU32 appears young except right after the birth, while OTU77 appears in the period right after the birth and in the second half of life.

Figs 3c and 3d show two examples of the original time series of OTUs with very small p-values for coexistence and exclusion, respectively. The light green shade indicates coexisting periods (1,1), the light orange indicates exclusive periods (1,0) or (0,1), and the white periods represent (0,0), which contribute to coexistence. Note that all of these examples are intuitively consistent as representatives of coexistence and exclusion. Conventional methods of data analysis may miss these examples because these time series contain a large number of zeros and appear to be non-stationary.

We calculate the p-values for all OTU pairs from all seven mice. For each OTU pair, we then evaluated the combined p-value using the p-values from all seven mice. There are numerous ways to define the combined p-value, and it is known that particular care should be taken when the number of data points is very different or when the characteristics of the data are different [33]. In this study, we use the most popular standard Fisher’s method, since the data sizes and observation conditions were nearly the same for the seven mice [34]. Let p_1_, p_2_, …, p_7_ be the p-values of the seven mice. We introduced the following test statistic, *S*, and estimated the combined p-value from the χ^2^ distribution with degrees of freedom 2×7=14:

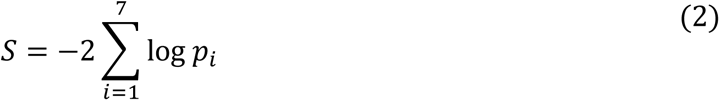

For each OTU pair, we compute the combined p-values for coexistence and exclusion separately based on this method. Fig 4a shows the cumulative distribution of the combined p-values for coexistence and exclusion, compared to the theoretical distribution of the null model (gray line). We set the threshold of statistical significance for the p-value at 10^-5^ (dotted line), consistent with the threshold p-value for a single mouse. The number of OTU pairs whose combined p-value for coexistence is less than this threshold is 18597, approximately 21.1% of the total pairs, while that for exclusion is 2347, approximately 2.7%. Interestingly, 18 OTU pairs show significance for both coexistence and exclusion individually, categorized as marginal.

**Fig 4.**
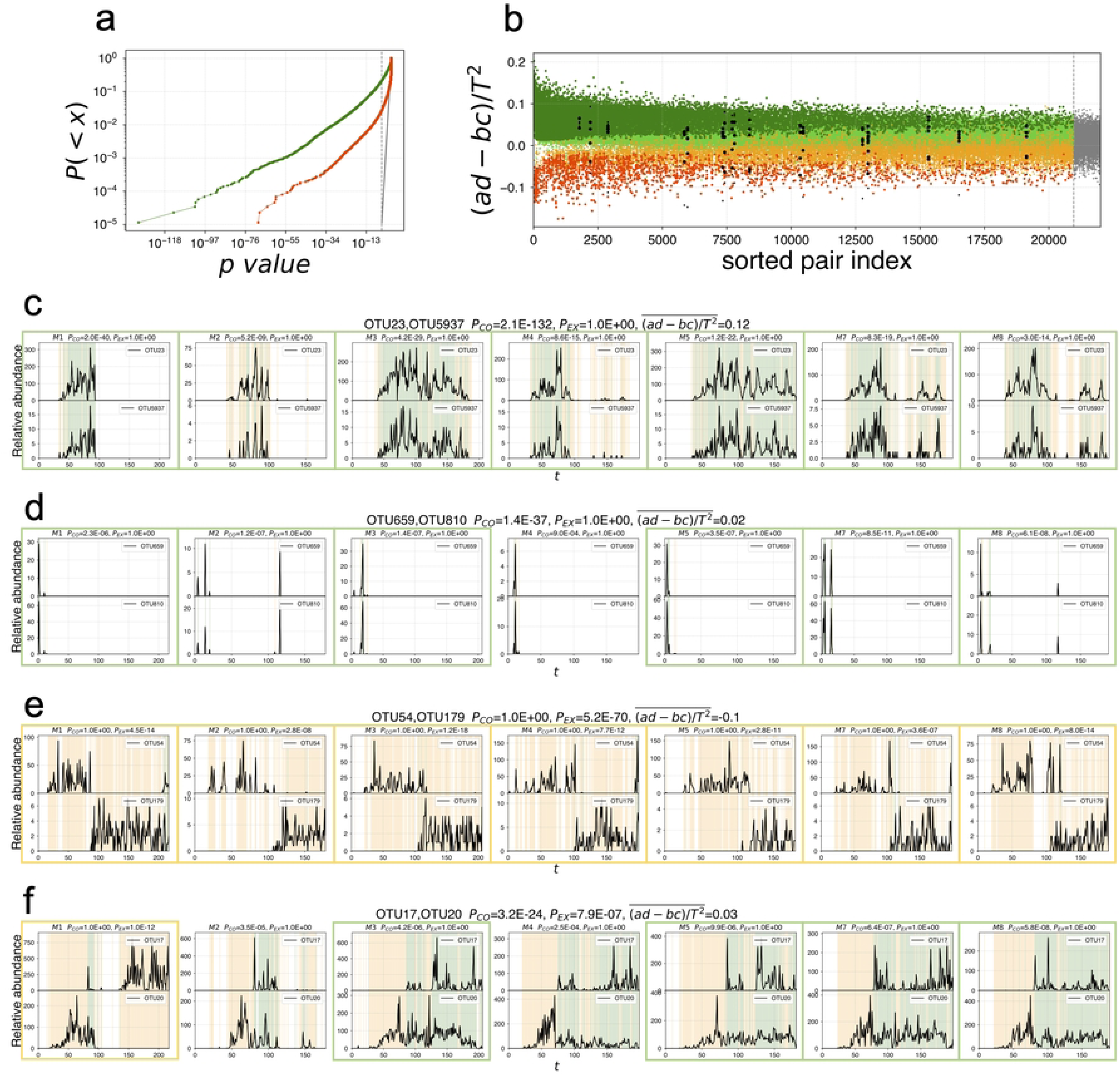
Coexistence and exclusion for seven mice. Fig 4a. Cumulative distribution of the p-values combined for all mice for coexistence (green) and exclusion (orange). The distribution of p-values in the case of the null model follows the same uniform distribution (gray) as the case of Fig 3a for one mouse. Fig 4b. Characteristics of significant coexistence and exclusion pairs. The vertical axis shows the value of polarity, *(ad-bc)/T^2^*; the horizontal axis represents the OTU pairs sorted in the ascending order of the combined p-value for the seven mice. For each OTU pair, 7 points are plotted corresponding to each mouse. Green dots are for significant coexistence pairs, dark green dots represent individually significant cases, and light green dots show individually non-significant cases. Orange dots are for significant exclusion pairs; dark orange and light orange represent the same meaning as green. Black dots show the marginal cases, larger dots show individually significant cases, and small dots show non-significant cases. Fig 4c. Examples of OTU pairs of time series for coexistence for seven mice. Green and orange shades show coexistence (1,1) and exclusion (1,0) or (0,1), respectively, while (0,0) are in white. Individually significant cases are framed in green for coexistence. Fig 4d. Examples of OTU pairs of time series for coexistence for seven mice with very small abundance. Individually significant cases are framed in green for coexistence. Fig 4e. Examples of OTU pairs of time series for exclusion for seven mice. Individually significant cases are framed in orange for exclusion. Fig 4f. Examples of OTU pairs of time series of the marginal case for seven mice. Individually significant cases are framed in green for coexistence and in orange for exclusion.

For all significant OTU pairs, we arranged them in ascending order of p-values on the horizontal axis and plot the values of (*ad-bc*)/*T^2^* for seven mice on the vertical axis as shown in Fig 4b. If the coexistence p-value per mouse is significant, it is plotted as a large dark green dot; if it is not significant per mouse, it is plotted as a small light green dot. Similarly, if the relationship is exclusive, it is plotted in orange. If the relationship is marginal, individual values are plotted in black with larger dots for individually significant cases. In Fig 4c, a typical time series of significant OTU pairs of coexistence is shown for all seven mice. Fig 4d shows a time series of a typical significant coexistence OTU pair with small *a* for all mice, confirming simultaneous appearances. Fig 4e shows a typical case of exclusive OTU pair for all mice; there, the orange periods showing exclusion dominate for all mice. Fig 4f shows a case of a marginal OTU pair, where coexistence is significant for mice M3, M5, M7 and M8, and exclusion is significant for M1.

### Detection of synchronization and antisynchronization

For a pair of time series whose number of co-occurrences, i.e. the number of (1,1) in the previous subsection is not zero, we can introduce a statistical test for up-down synchronization. To do this, we introduce the ternary transformation from the time difference of the original integer time series *x_i_*(*t*) to *y_i_*(*t*):

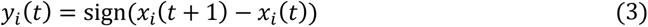

In order to quantify the strength of synchronization of OTU-*i* and OTU-*j* we introduce the following inner product *I*:

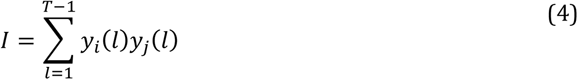

This quantity is positive when the up-down of the original time sequences are synchronized, and it is negative for an antisynchronized case. For this inner product, the corresponding p-value is estimated by comparing it with the null hypothesis model, in which the non-zero values of the time series {*x_i_*(*t*)} are randomly shuffled, while the points with *x_i_*(*t*)=0 are kept as 0. In this randomized time series, for the time point *t\** that fulfills *x_i_*(*t*\*)=0 and *x_i_*(*t*\*+1)=0, the value of *y_i_*(*t*\*) is always 0, and this time point does not contribute to *I*, so we neglect such time point in the calculation of *I*. For each pair of OTU-*i* and OTU-*j* the number of time points that can contribute to *I*, *L_ij_*, is counted as described in the Materials and Methods section, and the value of the inner product is normalized as *I*/*L_ij_*., which takes a value between −1 and 1. The corresponding p-value is calculated by a binomial distribution as explained in the Materials and Methods section. Fig 5a shows the results of the p-value distributions for synchronization (blue) and antisynchronization (red) for the mouse M1. The combined p-value distributions are plotted in Fig 5b, where the threshold of significance is the same as the former cases of coexistence and exclusion. We observe numerous significantly correlated OTU pairs; however, we should remove spurious correlations resulting from changes in compositional shares.

**Fig 5.**
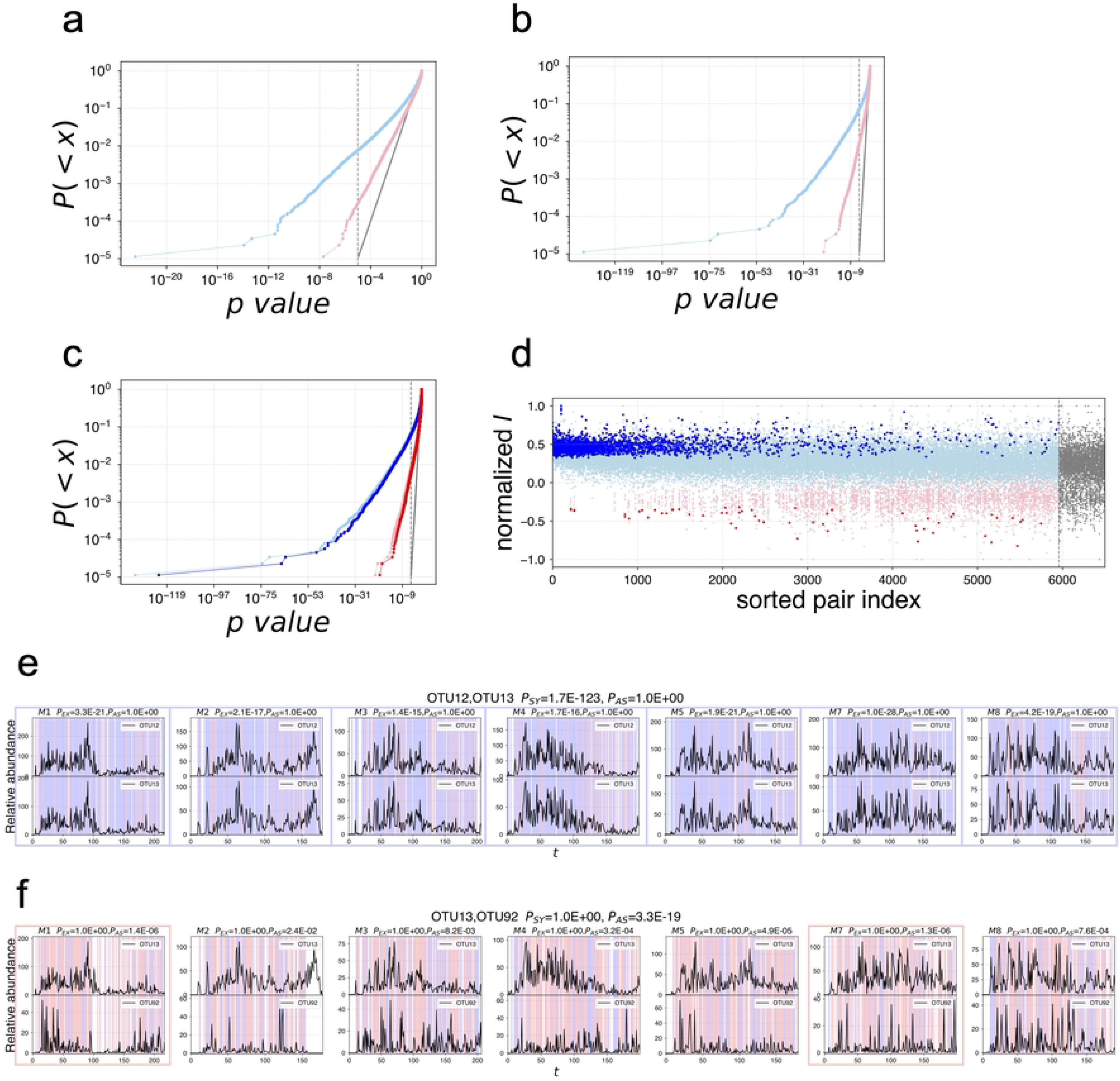
Synchronization and antisynchronization for seven mice. Fig 5a. Cumulative distribution of the p-values using the original time series for one mouse for synchronization (light blue) and antisynchronization (pink). The distribution of p-values in the case of the null model follows the same uniform distribution (gray) as the case of Fig 3a for one mouse. Fig 5b. Cumulative distribution of the combined p-values using the original time series for synchronization (light blue) and antisynchronization (pink). The distribution of p-values in the case of the null model follows the same uniform distribution (gray) as the case of Fig 3a for one mouse. Fig 5c. Cumulative distribution of the combined p-values after correction of spurious correlations for synchronization (blue) and antisynchronization (red) compared with the plots in Fig 5b. The distribution of p-values in the case of the null model follows the same uniform distribution (gray) as the case of Fig 3a for one mouse. Fig 5d. Characteristics of significant synchronization and antisynchronization pairs. The vertical axis shows the value of inner products, *I*, and, the horizontal axis represents the OTU pairs sorted in the ascending order of the combined p-value for the seven mice. For each OTU pair, 7 points are plotted corresponding to each mouse. Blue dots are for significant synchronization pairs, dark blue dots represent individually significant cases and light blue dots show individually non-significant cases. Red dots are for significant antisynchronization pairs. Dark red and pink depict the same meaning as blue. There is no marginal case. Fig 5e. Examples of OTU pairs of time series for synchronization for seven mice. Blue shades show synchronization (+1,+1) or (−1,−1), and red shades show antisynchronization (+1,−1) or (−1,+1), while the white shade shows the time points to be excluded for this analysis. Individually significant cases are framed in blue for synchronization. Fig 5f. Examples of OTU pairs of time series for antisynchronization for seven mice. Individually significant cases are framed in red for antisynchronization.

We consider instances of spurious correlations where an OTU, let it be called OTU-*i*, has a dominant share that changes drastically over time. For example, if *x_i_*(*t*)=2000 at time t and *x_i_*(*t*+1)=1000, then the sum of all other OTUs at time *t* is 1000, while it becomes 2000 at time *t*+1 maintaining a total sum of OTU is always 3000. Let us assume that this population change is purely caused by this OTU-*i* independently of all other OTUs. In this case, OTU-*i* will have antisynchronous correlations with all other OTUs, as the compositional shares of these OTUs would drop to half their average values. Simultaneously, any pair of other OTUs would show synchronous correlations, as their populations decrease together. To correct these spurious correlations, we hypothetically assume a situation where OTU-*i* did not exist, and the total sum of the sampling number of OTUs, excluding OTU-*i* is adjusted to 3000. Details of this new correction method of spurious correlation are described in the Materials and Methods section. We applied this correction for all possible combinations of OTUs, and judged the statistical significance by the condition that the corresponding p-values are always lower than the threshold. By this correction, the number of significant pairs decreased from 6,140 to 5,447 for synchronization and from 740 to 511 for antisynchronization.

In Fig 5c, the resulting distributions of the corrected combined p-values for synchronization and antisynchronization are shown in darker colors compared with the cases of no correction in lighter colors. Fig 5d is plotted similarly to Fig 4b; namely, for all significant OTU pairs, we arrange OTU pairs in ascending order of p-values on the horizontal axis and plot the values of *I*/*L_ij_* for seven mice on the vertical axis. If the synchronous p-value per mouse is significant, it is plotted as a large dark blue dot; if it is not significant per mouse, it is plotted as a small light blue dot; similarly, if the relationship is antisynchronous, it is plotted in red. It should be noted that there is no marginal case in which both synchronization and antisynchronization are significant individually. As known from this plot, there are many individually significant cases in the synchronization analysis, while there are fewer individually significant antisynchronization cases. This tendency is understood by the property that the number of time points *L_ij_* is generally smaller in the cases of antisynchronization because it is similar to an exclusive relation, and it is challenging to attain small p-values individually. Typical examples of time series for synchronization and antisynchronization are shown in Figs 5e and 5f; in both cases, the properties of all seven mice look similar.

Our results for all 87,990 combinations of OTUs are summarized in Table 1. Rows show coexistence relationships, and columns show synchronization relationships. The numbers for coexistence (CO), exclusion (EX), synchronization (SY), and antisynchronization (AS) means the number of OTU pairs whose combined p-value is smaller than the significant level. The numbers in brackets are the expected numbers if the coexistence property and synchronization property are independent. For example, 4,436 OTU pairs, both CO and SY, are significant, which is about four times bigger than the mean expectation of an independent random case (1,151). Similarly, the number of significant OTUs in both EX and AS (that is 70), is more than five times larger. For the case of both EX and SY, there was no OTU pair, while the independent random case is expected to be 147. From these results, it is evident that CO and SY, as well as EX and AS, are highly correlated, and EX and SY are highly negatively correlated.

**Table 1.** Results of the significant pair numbers.

|  | **SY** | **AS** | **MA** | **RA** |  |
| --- | --- | --- | --- | --- | --- |
| **CO** | 4436<br>(1151) | 58<br>(111) | 0 | 14103 | 18597<br>(21.1%) |
| **EX** | 0<br>(147) | 70<br>(14) | 0 | 2277 | 2347<br>(2.7%) |
| **MA** | 0 | 0 | 0 | 18 | 18<br>(0.02%) |
| **RA** | 1011 | 383 | 0 | 65634 | 67028 |
|  | 5447<br>(6.2%) | 511<br>(0.6%) | 0 | 82032 | 87990 |

CO for coexistence, Ex for exclusion, SY for synchronization, AS for antisynchronization, MA for marginal, RA for random meaning not significant. The numbers in brackets are the mean numbers expected if the horizontal columns and vertical columns are independent. It is confirmed that CO and SY, EX, and AS are strongly correlated, while EX and SY, CO and AS are negatively correlated.

Approximately 25% of all 87,990 OTU pairs show a significant relationship in one of the four statistical tests. Furthermore, it has been established that nearly all 420 OTUs participate in at least one of such significant relationship, with only one OTU identified as not belonging to any significant relation to other OTUs. Our results underscore the prevailing strength of interactions among OTUs, even among those with low populations characterized by numerous zero values.

### Interaction among phyla

As described in the preceding subsections, there are approximately 25% of significant interactions between OTUs. A part of these interactions is shown in Fig 6a by a network diagram connecting significant OTU pairs with the colored lines of corresponding significant statistical tests compared to the gene phylogenetic trees with eight phyla. This intricate interaction diagram shows a tendency that the OTUs belonging to the same phylum tend to have more CO (green) and SY (blue) links, while OTUs between different phyla tend to have more EX (orange) and AS (red) links. In order to quantify these properties, we categorize the OTUs into eight phyla and count the number of significant OTU pairs between the phyla. Figs 6b, 6c, 6d and 6e show a combination of phyla in which the number of significant OTU pair numbers is statistically high for CO, EX, SY and AS, respectively, compared with the numbers of the null model of independent random cases. There are 20 combinations of phyla whose p-value is at a significant level, less than 0.01, and the results are summarized in Fig 6f by a network diagram. Regarding CO and SY as cooperative, and EX and AS as antagonistic relationships, there are two cooperative groups of phyla, {Firmicutes, Deferribactreses}, and {Bacteroidetes, Verrucomicrobia, Tenericutes, Actinobacteria, TM7, Proteobacteria}, which are connected by CO links. The Firmicutes group seems to be antagonistic to the Bacteroidetes group, as these groups are connected by EX and AS links. Within the Bacteroidetes group, Proteobacteria has a marginal relation as it is also antagonistically linked by EX and AY to Bacteroidetes and also linked by EX to TM7.

**Fig 6.**
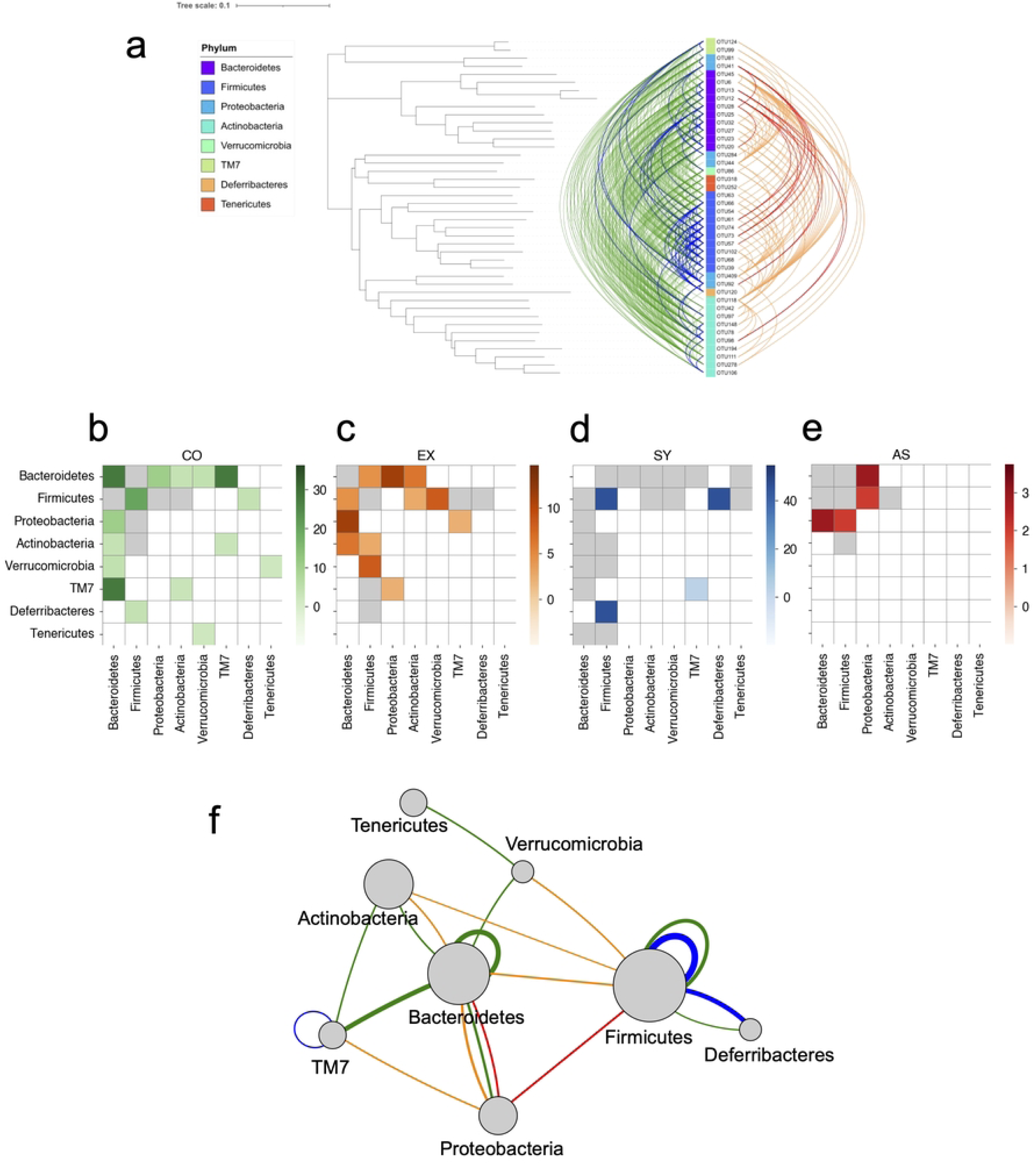
Interaction among eight phyla. Fig 6a. A part of significant relations between OTUs is categorized into 8 phyla with the gene phylogenetic trees. Significant pairs are connected by curved lines, CO (green), SY (blue), EX (orange), and AS (red). Fig 6b. Densities of significant CO between phyla. Darker green means more significant pairs than the null model, assuming random independent connection keeping the link numbers. The value in the green scale shows the absolute value of the logarithm of the p-value. Grey means the density is less than independent cases with the p-value less than 10^-2^. White represents the density, which is the level of independence. Fig 6c. The same plot as Fig 6b for EX is drawn in orange. Fig 6d. The same plot as Fig 6b for SY is drawn in blue. Fig 6e. The same plot as Fig 6b for AS is drawn in red. Fig 6f. Network diagram representing the relation between phyla. Green (CO) and blue (SY) lines show significant cooperative relations, and orange (EX) and red (AS) lines show significant antagonistic relations, with the thickness representing the statistical significance. The diameter of each node is proportional to the logarithm of the number of included OTUs.

## Discussion

In this paper, we introduced new data analysis methods that are designed to detect statistically significant correlations between any pair of OTU time series. As described in Sec.2.1, our data of OTU time series contain numerous zero values, no OTU has always existed in the seven mice for their whole lives; therefore, we thought that a careful treatment of zero values in the time series is key to detecting correlations among all OTUs.

To achieve this objective, in the “Detection of coexistence and exclusion pairs” subsection, we introduced the binary transformation (Eq (1)) for coexistence and exclusion, in which the data points of zero values play the central role. As shown in Fig 3a, we detected various significant OTU pairs for both coexistence (approximately 5%) and exclusion (approximately 1%) for the data of 1 mouse; additionally, by combining the results of seven mice, we found that 21% of pairs are significant in coexistence, and 2.7% in exclusion. As shown in Fig 4d, our method can detect significant coexistence for the cases with more than 97% of data points being zero values, and all seven mice showing a consistent property. Furthermore, as mentioned in “Detection of synchronization and antisynchronization” subsection, we confirmed that there is only 1 OTU that was not involved in significant coexistence nor exclusion pairs, and all other 419 OTUs have some significant correlations with other OTUs. We believe that the robust interactions among OTUs, even those with minimal presence, constitutes a noteworthy discovery. This outcome underscores the importance of directing greater attention toward OTUs with low populations to uncover the complex ecosystem of microbiomes.

In this study, to avoid inclusion by chance, the p-value threshold for the significance of the time series pairs was set to be 10^-5^, which is smaller than the inverse of the number of all pairs. This high standard was achieved for two reasons: one is the length of each time series that contains more than 200 data points, and the other is the parallel observation for seven mice. In fact, by combining the results of seven mice, the number of significant pairs increased three to four times for both coexistence and exclusion. In a case where the number of data points is half, namely approximately 100 data points, the p-values would become about square root of the original values; thus, it is roughly equivalent to make the threshold value to 10^-10^ in Fig 4a. We can estimate that the number of significant pairs will become approximately one-third. If the data points are about 50 and if we have only one mouse data, then we would be able to detect nearly 100 coexistence pairs and less than 10 exclusion pairs estimated from Fig 3a, assuming an imaginary threshold of 10^-20^. If the number of time points is less than 25, which corresponds to an imaginary threshold of 10^-40^, it would be difficult to detect significant coexistence or exclusion from 1 mouse data; however, by combining seven mice data, we will be able to detect more than 100 of significant pairs, as estimated from Fig 4a. It is important to prepare parallel experiments in the case there is a limitation in the number of data points.

In order to detect synchronization and antisynchronization, we introduced a null model that is created by randomly shuffling the values of raw data for non-zero time points, and we calculated the inner product values of the ternary transformed time difference sequence, representing up-0-down properties. It should be noted that we did not shuffle the data points of 0, so these analyses purely count the co-occurrence of ups and downs between the pair of OTUs. Compared with coexistence and exclusion analysis, the resulting p-values for synchronization and antisynchronization are much larger, meaning they are less significant. This is a natural consequence of the fact that the number of effective data points for OTU-*i* and OTU-*j*, *L_ij_*, is smaller, especially for antisynchronization. Both exclusion and antisynchronization are likely typical antagonistic relationships, and the number of *L_ij_*, becomes smaller for significant exclusion cases. In fact, the number of significant OTU pairs belonging to both exclusion and antisynchronization in Table 1 is 70, which is five times more than the number expected in the uncorrelated and random case (14). In this table, it should be noted that the number of OTU pairs that are significant in both exclusion and synchronization is 0, while the expected number is 147 in the uncorrelated and random case. These results are consistent with the assumption that both exclusion and antisynchronization represent antagonistic interactions between OTUs.

In the analysis of synchronization and antisynchronization in the subsection “Detection of synchronization and antisynchronization”, we introduced a new correction method to check spurious correlations caused by the compositional nature of the data. Details are described in the Materials and Methods section. The aim is to hypothetically remove the OTU-*i* of attention and make a proportional adjustment by integerizing the remaining OTUs so that the sum of the remaining OTUs becomes the whole number 3,000 of sampling. This correction removes the effect of decreasing or increasing the number of remaining OTUs due to an increase or decrease in the OTU-*i* of interest. Thus, it corrects for false negative correlations between OTU-*i* and OTU-*j*, as well as false positive correlations for pairs other than OTU-*i*. In the actual calculation, when correcting the correlation between OTU-*j* and OTU-*k*, the correction is calculated for all *i* (except *j* and *k*) when *i* is hypothetically removed, and the p-value of the pair, OTU-*j* and OTU-*k*, is given by the largest among all p-values, including the case when nothing is removed. As summarized in the Materials and Methods section, this correction reduces the number of significant pairs by approximately 30%. However, we confirmed that most of the strong correlations are still significant and that the functional form of the distribution of p-values is not much affected, as shown in Fig 5c. A merit of our correction method is the transparency of each procedure. We can quantitatively check which OTUs are causing spurious correlations in the shares of other OTUs, as shown in Fig 7.

**Fig 7.**
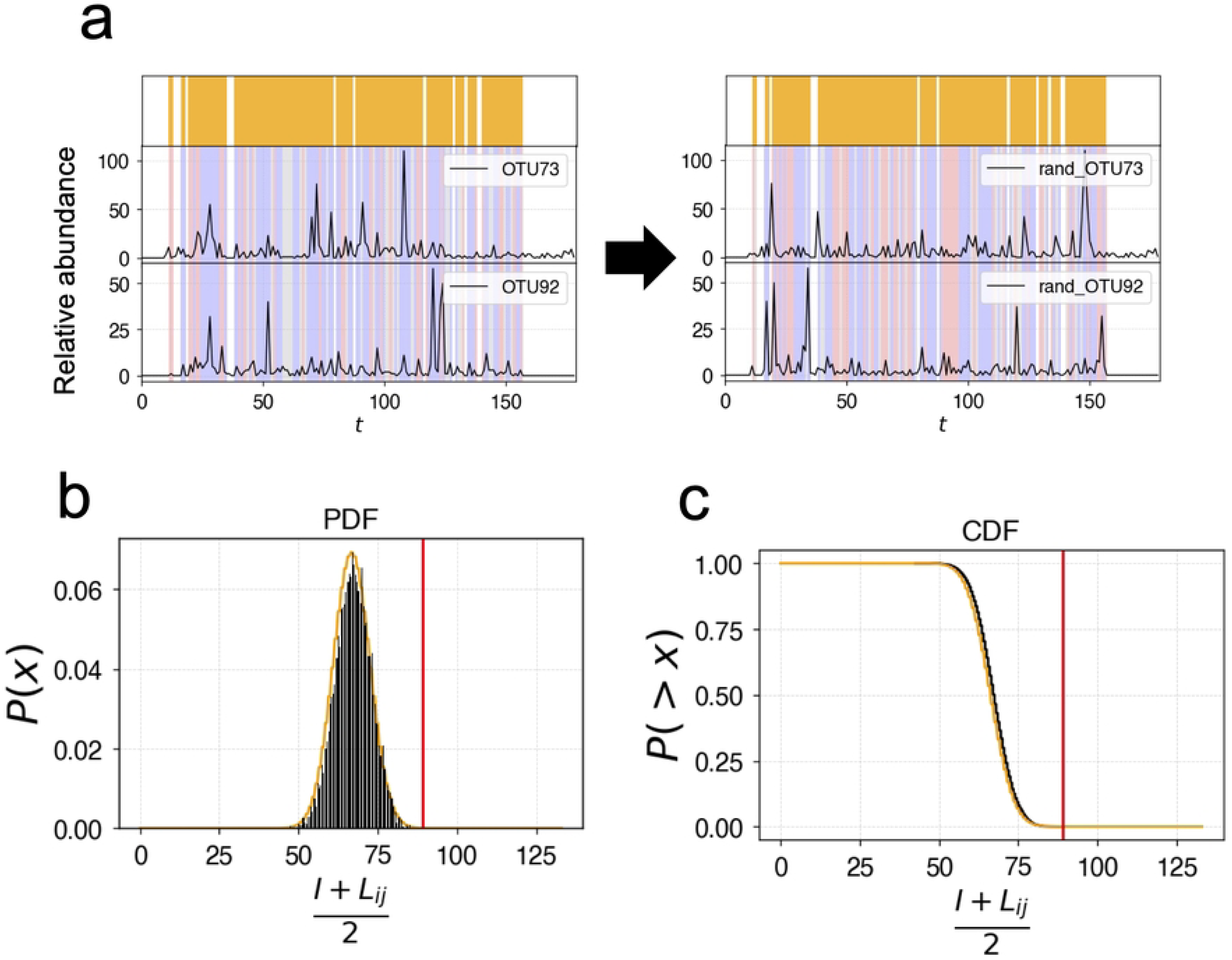
Random shuffling method for synchronization. Fig 7a. Randomly shuffled time series for calculation of synchronization. Left: The original time series of an OTU pair. Right: An example of a randomly shuffled time series, where shuffling is applied to non-zero points. The periods shown in yellow are the time points where synchronization is calculated, *L_ij_*. Synchronized time points are shaded in light blue, and antisynchronized points are shaded in pink, and the points that belong to neither are shaded in gray. Fig 7b. The probability density of the value, *(I+L_ij_)/2*. Black bars show the result of randomly shuffled null-models. The yellow curve presents the theoretical function approximately derived by the binomial distribution. The red line indicates the value for the real-time series. Fig. 7c. The cumulative distribution plot of Fig 7b. The p-value is estimated from the value of the cross point of the red line and the yellow curve.

In the “Interaction among phyla” subsection, we introduced the grouping of the OTUs into eight phyla and displayed the interactions between the phyla and themselves as a network diagram. It was confirmed that OTUs between the same phyla tend to be linked more cooperatively, while interactions with different phyla can be cooperative or antagonistic, as shown in Fig 6f.

In summary, the methods proposed in this paper are generally applicable to any similar data, such as integer-valued vector-type time series. We believe our methods can serve as basic general tools for the detection of statistically significant correlations. Note that if there is no regulation for the total sum of values, our four methods, CO, EX, SY and AS, can be used without the compositional correction. The methods CO and EX are suitable and powerful for data with numerous zero values. If the time series contains no or minimal number of zero values, then SY and AS will be useful to detect cooperative or antagonistic interactions. Our methods include no black-box and every detail of statistical tests can be checked directly by the p-values estimated by comparing them with the null models. The programming codes of these methods are available via GitHub.

## Materials and Methods

### The data

The data we used in this paper is the same data in reference [13] and the raw data is available from this reference. Initially, there were eight mice, M1 to M8; however, M6 died at a young age from cancer, and the others lived long and healthy for more than 820 days. In this paper, we omitted M6 and used the data for the other seven mice.

### Counting the numbers {*a,b,c,d*}

For the given time series for OTU-*i*, {*x_i_*(*t*)}, we introduce the transform, *s_i_*(*t*) = sign(*x_i_*(*t*)), which takes either 1 or 0 as {*x_i_*(*t*)} are non-negative integers. For the pair of OTU-*i* and OTU-*j*, the numbers {*a,b,c,d*} are calculated by the following forms of inner product.

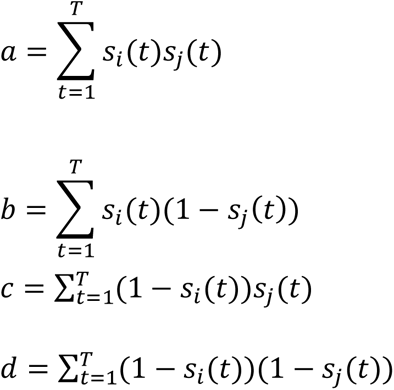

### Definition *Lij* and calculation of the p-value of up-down synchronization

In this subsection, we describe the definition of *L_ij_* for the pair of OTU-*i* and OTU-*j* and the way of calculation of p-value for the inner product *I* defined by Eq (4). The time points of *L_ij_* are those points that either *x*(t) or *x*(t+1) is not 0 for both *i* and *j*. It is calculated by the following equation.

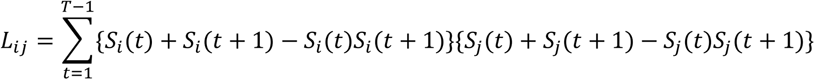

Next, we introduce the null model and estimate the p-value for the inner product *I*. The null model is defined by random shuffling of the non-zero values of the time series {*x_i_*(*t*)}, while the points with *x_i_*(*t*)=0 are kept as 0 as shown in Fig 7a. For a time series thus randomized {*x’*(*t*)}, we approximate that the value of *y’*(*t*)=sign{*x’*(*t*+1)-*x’*(*t*)} can be approximated by an independent random number +1 or −1 with probability 1/2. Then, the value of *I* can be approximated by the following equation:

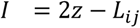

where z is a binomial random number taking a non-negative integer given by the probability density function, *B*(*L_ij_*, 1/2). In Fig 7b, the probability density function for the null model created by 10,000 random samples is plotted with the theoretical functional form of the binomial distribution; both distributions fit nicely. From this theoretical function, we can calculate the p-value by integrating the probability that is more extreme, as shown in Fig 7c.

### Correction of spurious correlations in compositional time series

Herein, we introduce a new method of correction of spurious correlations in compositional time series. The correlations we pay attention are schematically shown in Fig. 8a. At time *t*, let us assume the case that the abundances of OTU-*i*, OTU-*j*, and OTU-*k* are *x_i_*(*t*)=2000, *x_j_*(*t*)=200, *x_k_*(*t*)=100, with the total sampling number of OTUs at *t* being always *N*=3000. We also assume the case that these OTUs are independent and OTU-*j* and OTU-*k* are stationary keeping the same absolute density all the time. In the case that OTU-*i* changes drastically such as *x_i_*(*t*+1)=1000, *x_i_*(*t*+2)=2000 and *x_i_*(*t*+3)=1000, then the sum of abundance of all other OTUs are 1000 at time *t*, 2000 at time *t*+1, 1000 at time *t*+2, and 2000 at time *t*+3. We can expect that *x_j_*(*t*+1)=400, *x_k_*(*t*+1)=200, *x_j_*(*t*+2)=200, *x_k_*(*t*+2)=100, *x_j_*(*t*+3)=400, *x_k_*(*t*+3)=200, as schematically shown in Fig. 8a Left.

**Fig 8.**
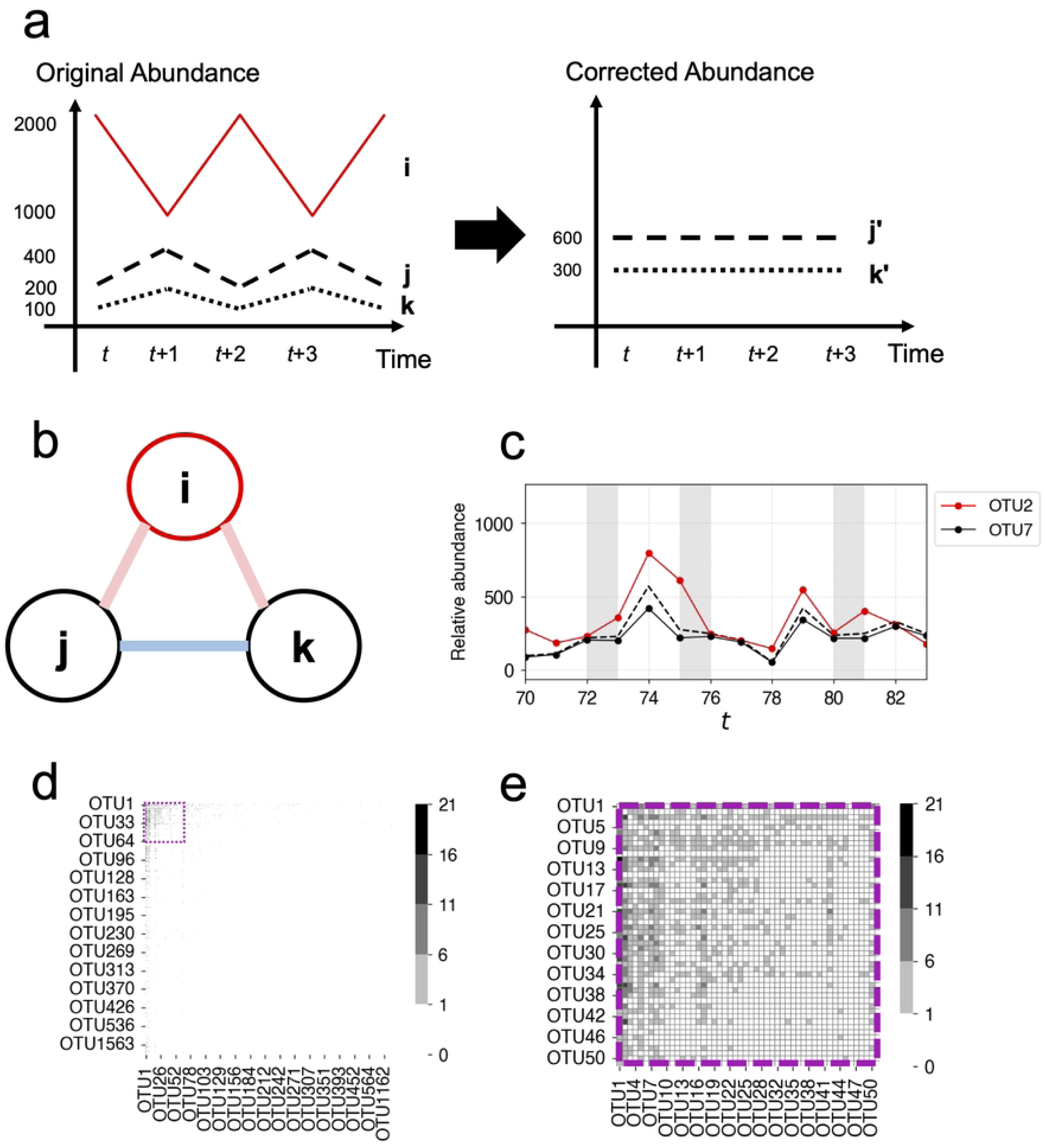
Correction of spurious correlations. Fig 8a. Schematic figure showing how to correct the spurious correlations. Left: OTU-*i* oscillates as {2000, 1000, 2000, 1000, 2000,…}, OTU-*j* oscillates as {200, 400, 200, 400, 200,…}, OTU-*k* oscillates as {100, 200, 100, 200, 100,…}. Right: OTU-*i* is imaginarily removed, and the rest of OTUs are normalized to make the sum to be 3000, then OTU-*j’* becomes flat as {600, 600, 600, 600, 600,…}, OTU-*k’* also becomes flat as {300, 300, 300, 300, 300,…}. Fig 8b. Schematic figure showing the spurious correlations. In the case of Fig. 8a Left, we can observe synchronization between OTU-*j* and OTU-*k*, and antisynchronization between OTU-*i* and OTU-*j*, and also OTU-*i* and OTU-*k*. These correlations vanish after correction, as shown in Fig 8a Right. Fig 8c. An example of correction for synchronization. OTU2 (red line) is imaginarily removed, and all other OTUs counts are corrected so that the portion of OTU2 eliminated is supplemented proportionally by other OTUs proportionally. The dashed line is the corrected time series of OTU7 with the original data shown by the black line. At the shaded periods, the up-0-down properties are changed by this correction. Fig 8d. Results of corrected numbers of up-0-down by the imaginary removal. The OTUs in the horizontal column are removed, and the counts of OTUs in the vertical column are corrected. This correction is effective only for OTUs with relatively high abundances, which are located in the left top area. The area surrounded by the purple dotted line is enlarged in the next figure. Fig 8e. Enlarged part of Fig. 8d. There are some OTUs that affects many other OTUs, and there are some OTUs that are affected by many other OTUs.

In order to correct these spurious fluctuations of OTU-*j* and OTU-*k* caused by the number change of OTU-*i*, we introduce an imaginary removal of OTU-*i*, namely, we select *N* samples without OTU-*i*. Then, the corrected numbers of OTU-*j* would be *x_j_*(*t*)= *x_j_*(*t+*1)= *x_j_*(*t+*2)= *x_j_*(*t+*3)=600. Those of OTU-*k* would be *x_k_*(*t*)= *x_k_*(*t+*1)= *x_k_*(*t+*2)= *x_k_*(*t+*3)=300, and no correlation would exist between OTU-*i* and corrected OTU-*j*, also between corrected OTU-*j* and corrected OTU-*k*. In this way the spurious negative correlation between OTU-*i* and OTU-*j*, OTU-*i* and OTU-*k*, and the spurious positive correlation between OTU-*j* and OTU-*k*, shown in Fig. 8b, can be corrected. This process of imaginary removal of OTU-*i* and the corrected value of OTU-*j* at time *t*, *x*^′^ (*t*), can be given by the following formulation.

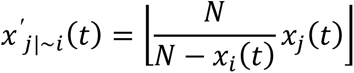

where ⌊*x*⌋ denotes the greatest integer function of a real number *x*. In Fig 8c, an example of this correction is shown for OTU-2 to be removed and OTU-7 to be corrected. The dotted line shows the corrected values of OTU-7, and those time points are shaded where the signs of up-0-down changed by this correction. We apply this imaginary removal process for all OTUs one by one, and calculate the correction for all other OTU time sequences. Fig. 8d and its partial enlargement, Fig. 8e, show how this imaginary removal of OTU-*i* affects OTU-*j* by counting the corrected numbers of signs in the time series of OTU-*j*. The horizontal axis shows the name of the imaginarily removed OTU and the vertical axis show the name of the corrected OTU, and the color of each column indicates the number of corrected up-0-down signs. The maximum number of changes is less than 20, and some OTUs are very influential to other OTUs; however, many small population OTUs do not affect other OTUs at all. The final result of the corrected p-value for OTU-*j* and OUT-*k* is given by the largest p-value among all corrected p-values by assuming the removal of OTU-*i* for all *i*, including the p-value estimated for the original time series without the correction. Namely, the significant cases are the cases in which all these p-values are less than 10^-5^. By this correction, the number of significant pairs of synchronization and antisynchronization decreases approximately 30%.

It should be noted that the sum of OTUs after correction is not exactly 3000, as the total number may be reduced by the fraction that is rounded down when the resulting number of corrections is converted to an integer. This method of imaginary removal and correction can be generalized to the removal of two or more OTUs. However, the number of combinations to be removed would be so large that the computational cost would diverge, so here the number of imaginary removals of OTU is limited to one.

## Code Availability Statement

All data and Python scripts used to perform this data analysis are available on GitHub. https://github.com/rie-maskawa/CESA

## Acknowledgments

This work was supported by JST, CREST Grant Number JPMJCR22N3, Japan.

